# A partial Drp1 knockout improves autophagy flux independent of mitochondrial function

**DOI:** 10.1101/2023.06.29.547095

**Authors:** Rebecca Z. Fan, Carolina Sportelli, Yanhao Lai, Said Salehe, Jennifer R. Pinnell, Jason R. Richardson, Shouqing Luo, Kim Tieu

## Abstract

Dynamin-related protein 1 (Drp1) is typically known for its role in mitochondrial fission. A partial inhibition of this protein has been reported to be protective in experimental models of neurodegenerative diseases. The protective mechanism has been attributed primarily to improved mitochondrial function. Herein, we provide evidence showing that a partial Drp1-knockout improves autophagy flux independent of mitochondria. First, we characterized in cell and animal models that at low non-toxic concentrations, manganese (Mn), which causes parkinsonian-like symptoms in humans, impaired autophagy flux but not mitochondrial function and morphology. Furthermore, nigral dopaminergic neurons were more sensitive than their neighbouring GABAergic counterparts. Second, in cells with a partial Drp1-knockdown and Drp1^+/-^ mice, autophagy impairment induced by Mn was significantly attenuated. This study demonstrates that autophagy is a more vulnerable target than mitochondria to Mn toxicity. Furthermore, improving autophagy flux is a separate mechanism conferred by Drp1 inhibition independent of mitochondrial fission.

## Introduction

Dynamin-related protein 1 (Drp1) is a member of the dynamin GTPase superfamily. Mutations in Drp1 cause severe neurologic phenotypes in humans^1^. This protein has four functional domains^2^. The N-terminal GTPase domain is essential for GTP hydrolysis. The middle domain is important for Drp1 self-assembly into dimers, tetramers, and higher-order oligomeric structures. The C-terminal GTPase-effector (GED) domain is important for mediating both intra- and intermolecular interactions of Drp1 proteins. The function of the small insert variable or B domain has not been fully resolved, but it has been reported to have lipid-binding region^3^, which is consistent with the observation that Drp1 binds to the outer mitochondrial membrane cardiolipin under stress conditions to induce mitochondrial fragmentation^4^. Overall, Drp1 is best known for its role in mitochondrial fission^5^. Upon activation in mammalian cells, Drp1 translocates from the cytosol to the outer mitochondrial membrane, where it binds to mitochondrial fission factor (Mff) and mitochondrial division proteins 49 and 51 (MiD49 / MiD51). Drp1 self-assembles into spiral structures which constrict and divide the mitochondrion into two daughter organelles^5^.

Although the role of Drp1 in mitochondrial division is critical to normal cellular function, excessive mitochondrial fission is detrimental to cells. Accumulating evidence indicates that partial Drp1 inhibition is protective in experimental models of neurodegenerative diseases such as Parkinson’s disease (PD), Alzheimer’s disease (AD), Huntington’s disease (HD), and Amyotrophic Lateral Sclerosis (ALS)^6^. The protective mechanism of blocking Drp1 function in these studies has been largely attributed to restoring the balance in mitochondrial fission and fusion. However, this conclusion may need to be re-considered, at least in some disease models. Despite its well-documented role in mitochondrial fission, Drp1 is mostly cytoplasmic. Under normal conditions, only approximately 3% of total Drp1 is cofractionated with mitochondria^7^. Live-cell imaging of fluorescently labelled Drp1 reveals that much of Drp1 appears diffuse in the cytoplasm and only approximately 2.5% of the total Drp1 puncta engage in mitochondrial fission^8^. Such observations, combined with the reports that Drp1 is also colocalized with other organelles such as lysosomes^9, 10^, endoplasmic reticulum^11^ and peroxisomes^12^, suggest that Drp1 most likely is a multifunctional protein that exerts its effects beyond mitochondrial fission. Indeed, Drp1 inhibition has been reported by independent laboratories to reduce protein aggregation in experimental models of PD^13, 14^, AD^15, 16^ and HD^17^, indicating the potential involvement of protein removal pathways such as autophagy. However, because these models impair mitochondrial function, which may result in impaired autophagy, it is not feasible to untangle with certainty whether Drp1 inhibition reduces protein aggregation via mitochondria, autophagy or a combination of both. A model with impaired autophagy independent of mitochondrial involvement is needed to evaluate this potential novel protective mechanism of Drp1 inhibition.

Manganese (Mn) is an essential trace metal required for normal development and function because it is essential for the activities of a variety of enzymes, including glutamine synthetase and hydrolases, and it is a cofactor for a variety of metalloproteins, including the mitochondrial Mn-superoxide dismutase (MnSOD)^18–20^. However, excessive exposure to Mn can cause parkinsonian-like symptoms known as manganism. Although symptoms resemble those of PD, manganism and PD are two distinctive disorders^21–23^. However, both human and experimental data support the notion that chronic low-level Mn exposure represents a risk factor for developing PD, as well as accelerating the disease onset and progression of PD by interacting with an individual’s susceptible genes^24–27^. Indeed, higher blood levels of Mn has been reported in PD patients^28^. Experimentally, chronic exposure to Mn in rat results in increased Mn levels in dopamine (DA) neurons of the substantia nigra *pars compacta (*SNpc)^29^. In combination, evidence suggests that Mn may contribute to PD pathogenesis.

The link between Mn toxicity and mitochondrial dysfunction has been made since the early observations that at high concentrations, Mn enters mitochondria through the Ca^2+^ uniporter channel^30^ and accumulates in the mitochondrial matrix, resulting in impaired oxidative phosphorylation^31^. However, the negative impact of Mn on mitochondria at sublethal concentrations has been questioned in a recent study^32^. Furthermore, under conditions where Mn induces mitochondrial dysfunction, autophagy is also affected^33^. Given the bi-directional relationship between mitochondria and autophagy^34^, it is not clear whether mitochondrial dysfunction precedes autophagy blockade or vice versa for Mn neurotoxicity. Combined with the observation that Mn is accumulated to a greater extent in lysosomes than in mitochondria^10^, it is critical to investigate whether autophagy or mitochondria is the initial vulnerable target of Mn.

In this study we provide data to address two major gaps of knowledge in the field: First, by titrating the doses of Mn and by using multiple cell and animal models, it is evident that autophagy flux, not mitochondria, is the initial target of Mn. Second, using Mn as *in vitro* and *in vivo* models to impair autophagy without affecting mitochondria, a partial Drp1 knockout improved autophagy flux, indicating this protective mechanism is mediated independent of mitochondria.

## Results

### Mn impairs autophagy at low concentrations

To investigate whether mitochondria or autophagy is the initial vulnerable target to Mn, a systematic approach is needed to characterize the impact of Mn on mitochondria and autophagy flux. To this end, we started with cell culture models because they were more amenable to genetic and pharmacological manipulations. We performed time-course and dose-response cytotoxicity studies by exposing rat dopaminergic neuronal N27 cells, and HeLa autophagy reporter cells to increasing concentrations of MnCl_2_ (62.5 – 2000µM). Cell viability and Lethal Concentration-50 (LC_50_) values were calculated after 24h and 48h of MnCl_2_ treatment. Mn decreased cell viability in a time-dependent manner in both cell models (Fig. 1a,b) with LC_50_ values of 323.7 ± 8.7 µM (24h) and 314.5 ± 15.8 µM (48h) for HeLa cells, 260.6 ± 7.5 µM (24h) and 186.9 ± 2.5 µM (48h) for N27 cells. Next, using autophagy HeLa reporter cells that stably express mRFP-GFP-LC3 (Fig. 1c) to monitor autophagy flux^35^, we performed dose-response studies to determine the effects of Mn on autophagy (Fig. 1d-e). After Mn treatment for 24h, the number of autophagosomes increased, in a dose-dependent manner, while autolysosomes markedly decreased compared to control cells. These results are indicative of a blockade in autophagy flux. At our lowest tested Mn concentration (62.5 µM), autophagy flux was severely impaired.

We further investigated whether Mn impaired the early or late stage of autophagy flux. First, we assessed the late stage using chloroquine (CQ), a lysosome inhibitor that interferes with the function and fusion of autophagosomes with lysosomes^36^. We treated HeLa autophagy reporter cells in the presence or absence of MnCl_2_, CQ or a combination of both. We hypothesized that since CQ blocks downstream autophagy, in the scenario where Mn predominantly blocks early stage of autophagy (induction or autophagosome function), more autophagy blockade would occur due to a combination of blocking autophagy flux at two separate stages. As seen in Fig. 1f,g, CQ blocked autophagy flux in a dose-dependent manner (50 and 100µM). When combined with CQ 50µM, a concentration that did not maximally inhibit autophagy, Mn at the concentration of 125µM did not further impair autophagy – suggesting that Mn does not inhibit autophagy at an early stage, and as compared to CQ, it is a weaker inhibitor at the late stage. As a complementary approach, we assessed the levels of Atg5, a marker for early autophagy. We treated the dopaminergic neuronal N27 cells with Mn 125µM, with or without CQ 50µM. As seen in the immunoblotting data (Fig. h,i), a combination of CQ and Mn did not further increase the levels of Atg5 – further supporting the lack of effects of Mn at upstream autophagy. Together, these data demonstrate that at a sublethal concentration, Mn inhibits the late stage of autophagy.

**Fig 1.**
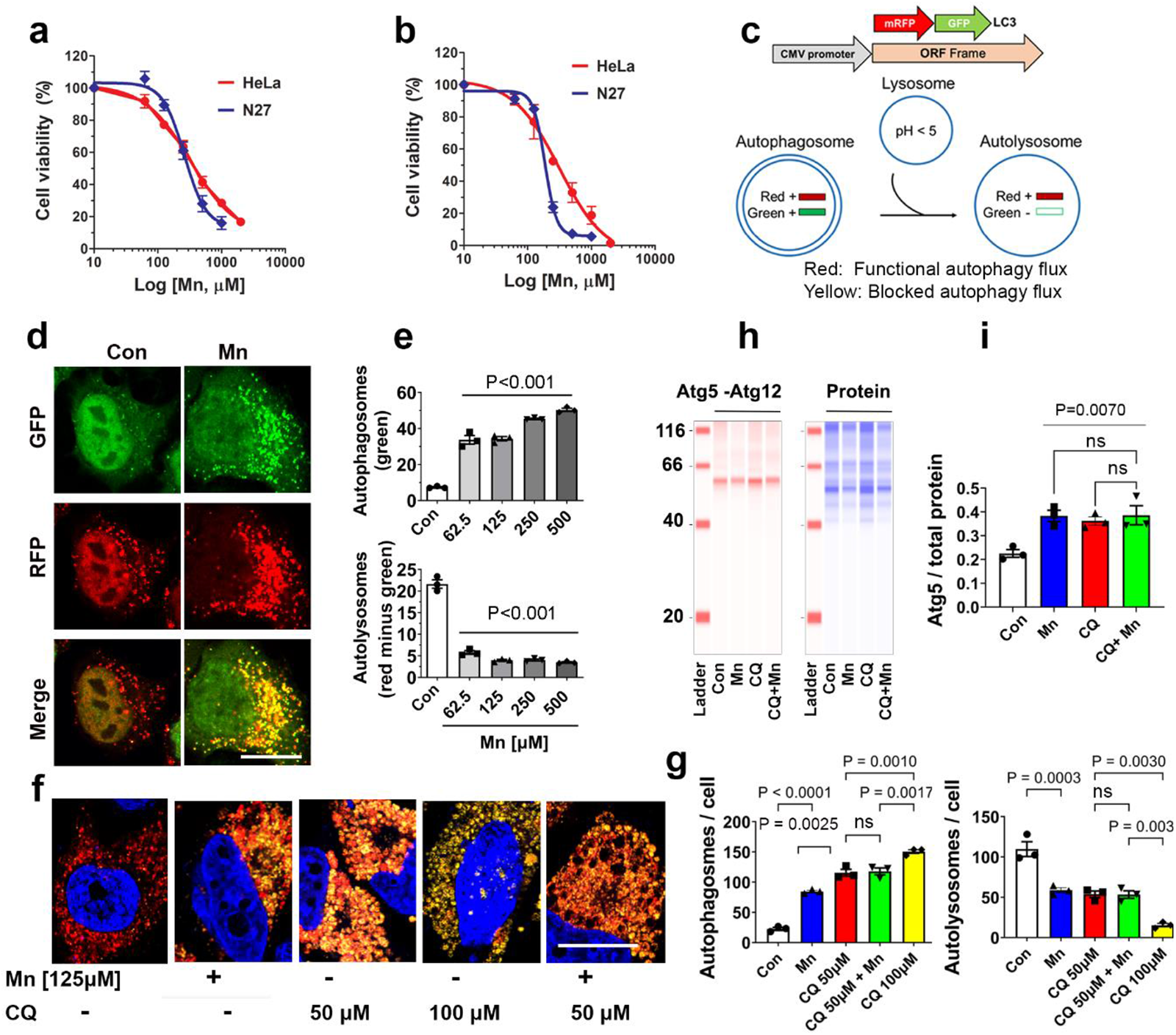
Dose-responses of Mn on cell viability and autophagy flux. **a**, Stable HeLa cells overexpressing mRFP-GFP-LC3, and the rat dopaminergic neuronal cells N27 were exposed to MnCl_2_ (10µM to 2mM) for 24h, or **b,** 48h. Cell viability was evaluated using Calcein AM dye. Data represent mean ± SEM, n=3 independent experiments with 4 wells per experiment. **c,** Schematic diagram illustrating the autophagy flux pathway and the construct used to create the mRFP-GFP-LC3 stable HeLa reporter cells. With this cell model, autophagosomes appear yellow due to the colocalization of RFP and GFP signals. Red signal indicates the flux is functional because the green signal is quenched by the acidic environment of the lysosomes, which fuse with autophagosomes. These stable cells were treated with vehicle control or Mn (62.5 – 500μM) for 24h. **d,** Representative images of cells treated with Mn (62.5µM) were captured using confocal microscopy. **e,** Green and red vesicles per cell were quantified using Fiji. Green vesicles represent autophagosomes. The number of autolysosomes was calculated by subtracting the number of green from the red puncta per cell. Scale bar = 20μm. **f,** Stable HeLa cells were treated with MnCl_2_ (125µM), chloroquine (CQ) or both. **g,** The number of autophagosomes and autolysosomes were quantified. Data represent mean ± SEM. **h,** N27 cells were treated with Mn 125 µM, CQ 50 µM or both for 24h. Cells were then collected and immunoblotted for Atg5 using Jess (ProteinSimple, Inc.). **i,** Total protein per capillary was used as loading control to normalize the levels of Atg5. All data represent mean ± SEM (n=3-4 independent experiments with approximately 30 cells per experiment for **e** and **g)** and groups were compared by using Kruskal-Wallis ANOVA test with Dunn’s post-hoc test; ns: non-significant.

### Mn does not affect mitochondria at the low concentrations that impair autophagy

To investigate whether at the low concentrations that Mn impaired autophagy flux (as seen in Fig. 1) mitochondria would also be impacted, we assessed mitochondrial morphology and function in the same cell model. To this end, a dose-response study of Mn (62.5 µM-250µM) was performed in HeLa autophagy reporter cells. Mitochondrial morphology was visualized using TOM-20 immunostaining followed by confocal microscopy (Fig. 2a). To quantify mitochondrial morphology and network more objectively, we used Mitochondrial Network Analysis (MiNA) of Fiji Image J. As shown in Fig. 2b, Mn did not cause structural abnormality at 62.5µM (data not shown) and 125 µM – a concentration well above the inhibitory effect of Mn on autophagy flux. At a higher concentration (250µM), Mn induced mitochondrial fragmentation. To further validate the effect of Mn on mitochondria, Seahorse extracellular flux analyzer was used to measure mitochondrial respiration. Consistent with the morphological data, mitochondrial function was not impaired by Mn unless it reached 250µM (Fig. 2c). Together, these morphological and functional studies of mitochondria indicate that autophagy flux is more vulnerable than mitochondria to Mn toxicity in this cell model.

### Drp1 inhibition attenuates impaired autophagy induced by Mn in Hela autophagy reporter cells

To evaluate the effects of blocking Drp1 on autophagy flux, HeLa autophagy reporter cells were transfected with siRNA-Drp1 for 24h and then treated with or without MnCl_2_ (62.5µM) for another 24h (Fig. 2d). We previously validated that this siRNA-Drp1 achieved approximately 70-80% of Drp1 knockdown efficiency in the cell types used in this study, whereas scramble-siRNA did not affect Drp1 levels^14^. As shown in Fig. 2e, Drp1 knockdown significantly attenuated autophagy blockade induced by Mn as evidenced by a reduction in the number of autophagosomes (from 41.35 ± 0.91 to 27.21 ± 2.08) and increased autolysosomes (from 3.73 ± 0.66 to 16.84 ± 0.94), respectively. In combination, these data indicate that reduced Drp1 improves autophagy flux in a cell model with impaired autophagy independent of abnormal mitochondrial function and morphology.

**Fig 2.**
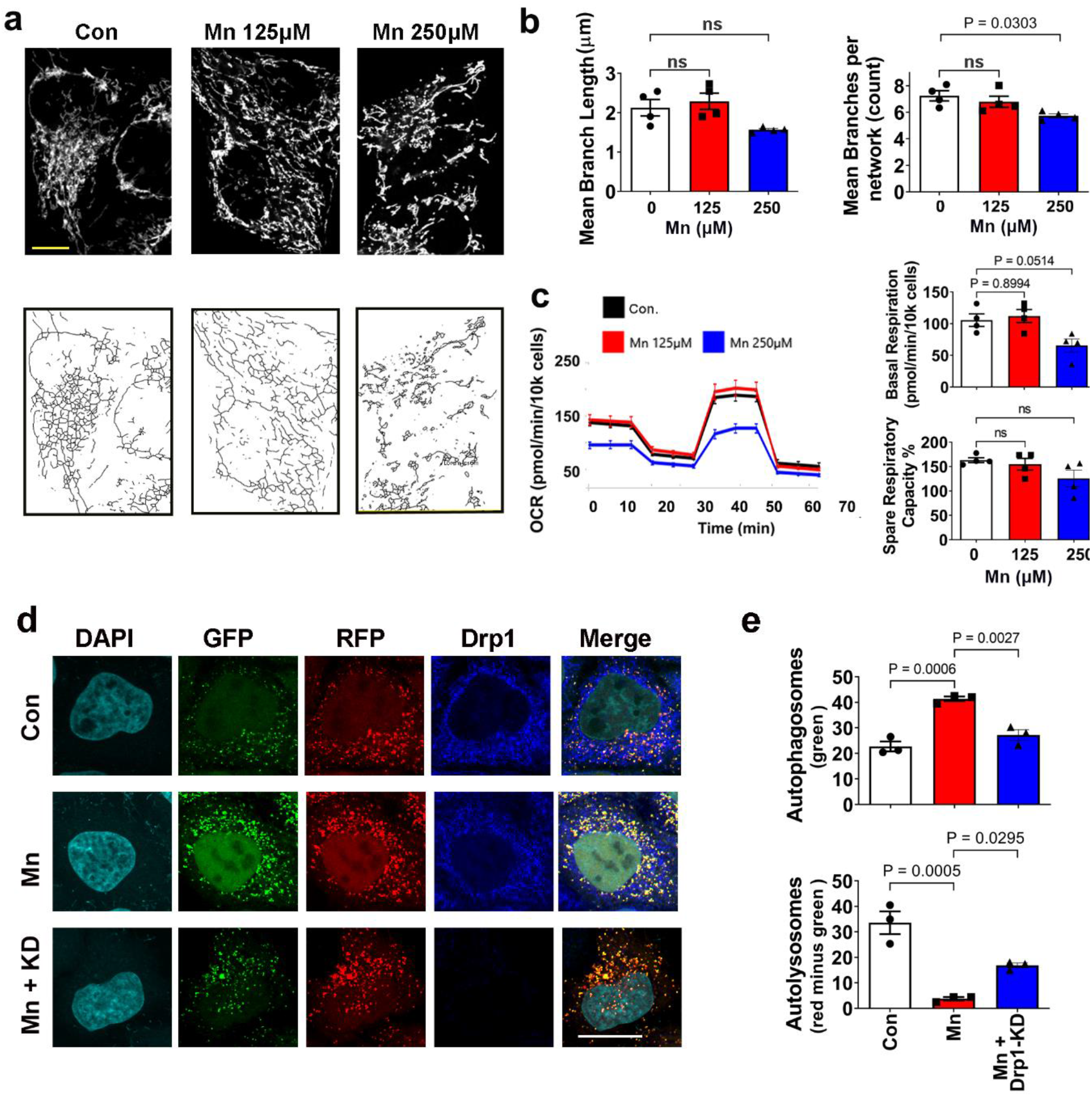
Low Mn doses do not affect mitochondria and inhibition of Drp1 attenuates its blockade on autophagy. HeLa autophagy reporter cells were treated with vehicle (DMEM media), 62.5μM (not shown), 125μM or 250μM of Mn for 24h. **a,** Representative confocal images of mitochondrial morphology after TOM20 immunostaining (upper panels, scale bar 20 μm), and then skeletonized (lower panels) for subsequent analysis of mitochondrial morphology. **b,** The MiNA plugin of Fiji was used to quantified mitochondrial morphology/network. Data represent the mean ± SEM (n=4 independent experiments), analyzed by one-way ANOVA, followed by Tukey’s post hoc test. **c,** Mitochondrial respiration was assessed by measuring OCR using the XFe96 Extracellular Flux Analyzer. Data represent mean ± SEM, n=4 independent experiments. **d,** HeLa autophagy reporter cells were transfected with siRNA-Drp1 for 24h to knockdown (KD) Drp1, and then treated with Mn (62.5μM) for another 24h, followed by immunostaining for Drp1. **e,** Images were captured and the number of autophagosomes and autolysosomes were quantified as described in Fig. 1. Data represent mean ± SEM (n=3 independent experiments with approximately 30 cells per experiment) and groups were compared by using Kruskal-Wallis ANOVA test with Dunn’s post-hoc test. Scale bar = 20μm.

### Drp1 inhibition attenuates impaired autophagy and protein aggregation induced by Mn in dopaminergic neuronal cells

To further corroborate the observations in HeLa cells as shown in Fig. 1 & 2, we used the N27 rat dopaminergic neuronal cells with ecdysone-inducible expression of human wild-type α-synuclein^14^. First, we assessed the effects of Mn on autophagy flux and whether reduced Drp1 levels would also be protective in these neuronal cells by quantifying the number of LC3 and p62 immuno-positive puncta per cell. As shown in Fig. 3a,b, Mn (125μM) alone increased both LC3 (39.72 ± 0.76) and p62 (54.30 ± 4.73) puncta compared to vehicle cells (LC3:18.06 ± 1.12; p62: 27.46 ± 0.99), indicating a blockade in the autophagy pathway. In contrast, inhibition of Drp1 attenuated the accumulation of LC3 (25.02 ± 2.06) and p62 (33.55 ± 2.36) puncta induced by Mn. These results are consistent with those in the HeLa cells (Fig. 2). In addition, upon co-treatment of ponasterone A (ponA, an ecdysone analog used to induce α-synuclein expression) and Mn, the number of LC3 (48.83 ± 3.54) and p62 (71.16 ± 8.33) puncta increased to levels that were statistically different to cells exposed to PonA (LC3, 33.23 ± 1.31; p62, 46 ± 2.20) and Mn alone (LC3, 39.72 ± 0.76; p62 54.30 ± 4.73). These results indicate that α-synuclein alone blocked autophagy, as did Mn, however a more pronounced inhibition was observed when α-synuclein was combined with Mn. Drp1 knockdown using siRNA significantly reduced LC3 and p62 puncta in both ponA induced cells (LC3, 21.43 ± 1.73; p62, 29.69 ± 1.78) and cells co-treated with ponA and Mn (LC3, 26.75 ± 0.5; p62, 33.16 ± 1.35), to a level that was not statistically different to vehicle-treated cells (LC3, 17.35 ± 0.71; p62, 25.57 ± 0.57).

Given that autophagy is a primary pathway by which misfolded/aggregated α-synuclein is removed, we assessed the effects of Mn and a partial Drp-1 knockdown on the levels of α-synuclein in this cell model. These cells accumulate proteinase-K resistance α-synuclein after two days of induction by ponA (vehicle control: 0.78 ± 0.31; ponA: 9.35 ± 0.40)(Fig. 3c,d). However, a non-toxic dose of Mn (125μM) significantly increased the number of these aggregates (19.46 ± 3.00). Knocking down Drp1 drastically reduced such protein aggregation (6.98 ± 0.75). Overall, these data suggest that autophagy dysfunction and α-synuclein aggregation induced by Mn-could be ameliorated by Drp1 inhibition.

**Fig 3.**
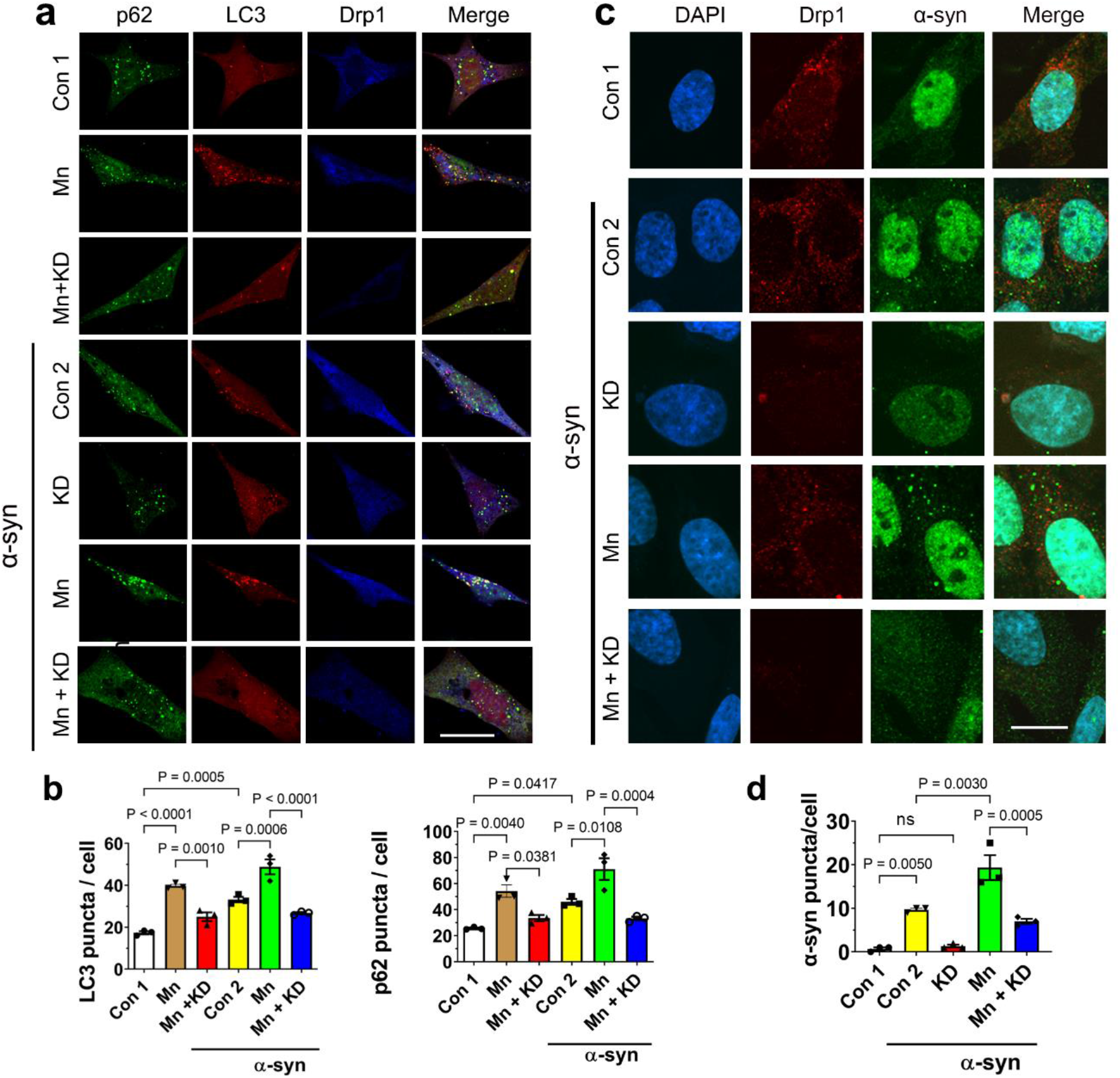
Drp1 inhibition protects against impaired autophagy and protein aggregation induced by Mn in dopaminergic neuronal cells. Stable N27 cells expressing inducible human wild-type α-synuclein were co-transfected with siRNA-Drp1 (KD) and LC3-cherry (due to low endogenous LC3 levels in this cell type) for 24h. **a,** Cells were then treated for another 24 h with MnCl2 (125μM) or vehicle controls (Con 1 & Con 2), in the presence or absence of PonA (20μM) to induce α-synuclein expression. **b**, Following confocal imaging, the number of green and red vesicles, representing respectively p62 and LC3 puncta was quantified using Fiji ImageJ. Data represent mean ± SEM (n=3 independent experiments, with at least 20 LC3 positive cells for each condition). Kruskal-Wallis ANOVA was used to compare between groups with Dunn’s post-hoc test. Scale bar 20μm. **c**, N27 cells were transfected with siRNA-Drp1 (10nM) for 24h, followed by α-synuclein overexpression induction (PonA 20μM), with and without Mn (125μM) for 48h hours. After post-fixation, cells were incubated with Proteinase K and then incubated with an antibody that detects α-synuclein (Millipore, AB5038). **d**, Fiji ImageJ was used to quantify α-synuclein-positive puncta per cell with a minimum of 30 cells per condition. Data are shown as mean ± SEM (n=3 independent experiments). Scale bar 20μm.

### Mn does not perturb mitochondrial morphology and function at low concentrations in dopaminergic neuronal cells

In the autophagy reporter HeLa cells, we observed that autophagy was inhibited by Mn at concentrations that did not affect mitochondria (Fig. 2). We asked whether Mn would also have the same effects in neuronal cells. Using N27 dopaminergic neuronal cells, we performed a dose-response study of Mn (62.5-250µM) and quantified mitochondrial morphology and function as done in Fig. 2. As shown in Fig. 4, Mn did not cause abnormal mitochondrial phenotype and function at 62.5 µM (data now shown) and 125 µM – concentrations that impaired autophagy flux in this cell type. At a higher concentration (250µM), Mn did induce mitochondrial fragmentation and impaired function.

**Fig 4.**
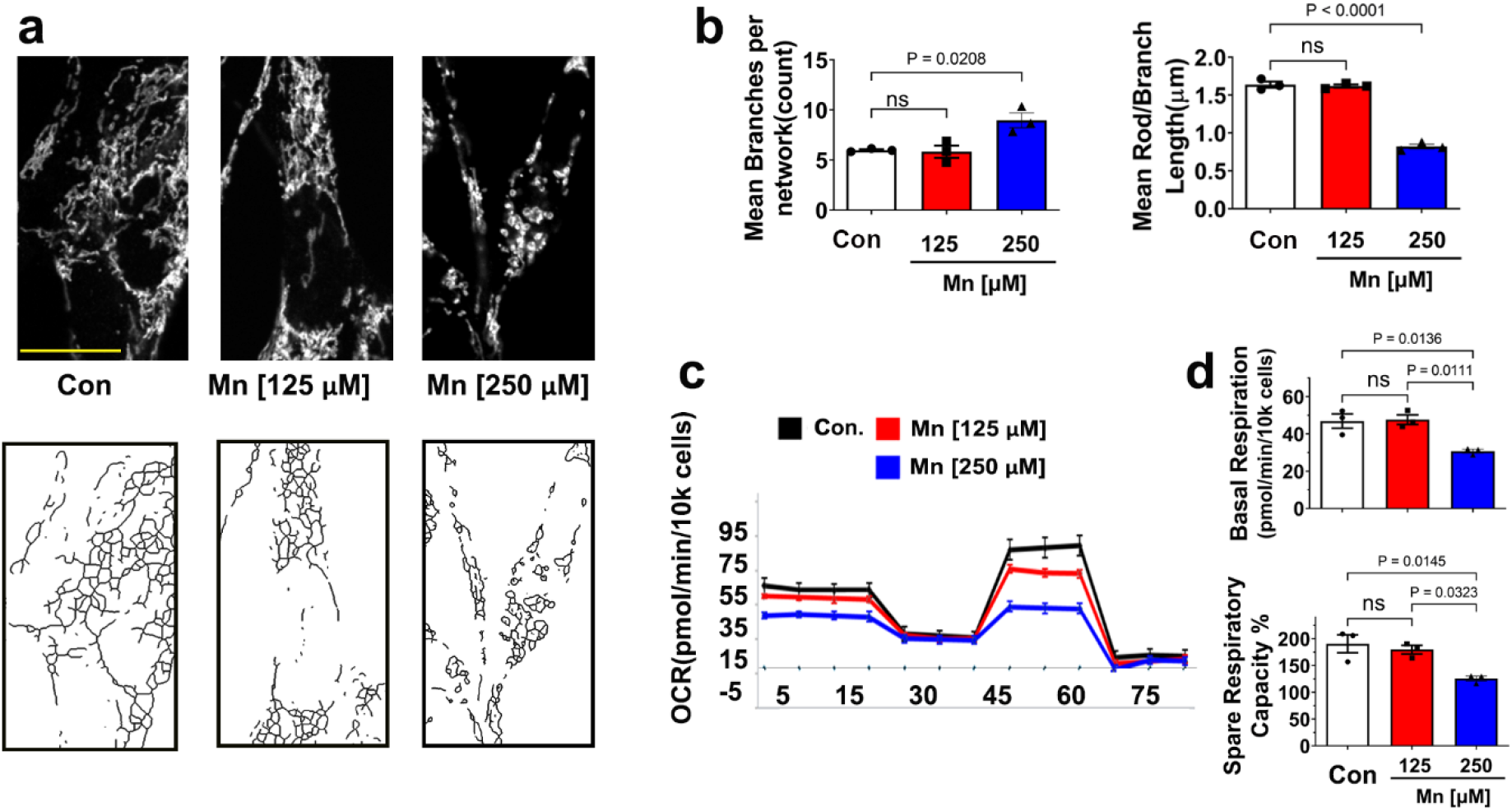
Mn does not affect mitochondria at low concentrations in dopaminergic neuronal cells. N27 neuronal cells were treated with 125μM or 250μM of MnCl2 or vehicle control (RPMI media) for 24h. **a,** Representative confocal images of mitochondrial morphology after TOM-20 immunostaining (upper panels), then skeletonized (lower panels) for subsequent analysis. Scale bar 20μm. **b,** Various parameters of mitochondrial morphology were quantified using Fiji MiNA plugin. Data represents mean ± SEM (n=3 independent experiments), analyzed by Kruskal-Wallis ANOVA test. **c, d,** Mitochondrial respiration was assessed by measuring OCR using the XFe96 Extracellular Flux Analyzer. Data represent mean ± SEM, n=3 independent experiments.

### Transcriptomic analysis reveals Mn impairs autophagy but not mitochondrial pathways

### in vivo

As demonstrated in our *in vitro* data using complementary approaches (Fig.1-4), when the concentration of Mn was titrated, it was clear that the inhibitory effects of Mn on autophagy and mitochondria were dose-dependent. To investigate the effects of Mn on autophagy and mitochondria *in vivo*, we started with an unbiased approach by performing RNAseq to identify changes in the transcriptomic profile in the ventral middle brain of the C57/BL6 mice treated orally with either MnCl_2_ (15mg/kg/day) or water control once daily for 30 consecutive days^37^. This Mn regimen was selected because it had been shown to induce dopaminergic neurodegeneration, protein aggregation and exosome release in mice with viral-mediated α-synuclein overexpressing^37^. We found 508 genes were up-regulated and 509 genes were down-regulated in response to Mn exposure (p<0.05) (Fig. 5a). Among those differentially expressed genes (DEGs), we further profiled the ones that were specifically involved in the autophagy pathways using “Autophagy Transcription Gene Toolbox”^38^. Results revealed that the four subsets of autophagy genes that were significantly dysregulated by Mn treatment belong to “mTOR and upstream pathways”, “Autophagy Core”, “Autophagy Regulators” and “Lysosome” (*p*<0.05) (Fig. 5b). No sex differences were observed within the Mn-treated mice as well as the control mice; therefore, we combined the results (n = 4 females and n = 4 males per group) for these autophagy associated DEGs (Supplementary File 1). Because some of these genes have either inhibitory or activating effects on autophagy function, it is not feasible to conclude whether or how these alterations would ultimately affect autophagy. To this end, we used KEGG pathway analysis to examine whether the autophagy pathways (Supplementary Fig. 1) would be enriched either in “up-regulated” or “down-regulated” pathways after exposure to Mn. The results showed a significant (padj=0.0128) downregulation in the Mn-treated mice as compared to the vehicle control group (Supplementary File 1, Supplementary Fig-1). These results indicate the impairment of autophagy upon Mn exposure *in vivo*.

To assess the effects of Mn on mitochondria in these animals, genes that are associated with mitochondrial dynamics and mitochondrial function were compared between the Mn-treated mice and control mice. As shown in Fig. 5c, no genes involved in mitochondrial dynamics were significantly affected by Mn treatment, and 97.7% (128/131) genes responsible for mitochondrial function were not affected (Fig. 5d, Supplementary File 1). The three significantly affected genes were mt-Nd4 (P=0.01845, Log2Fold Change=0.1405), mt-Nd1 (P=0.04453, Log2Fold Change=0.1382) and Sdhb (P=0.04861, Log2Fold Change=0.1841), all of which are subunits of mitochondrial complex 1 and slightly upregulated after exposure to Mn (Fig. 5d, Supplementary File 1). Cumulatively, these results indicate that the repeated low-dose Mn treatment impairs autophagy but not the pathways related to mitochondrial dynamics and mitochondrial function. These *in vivo* data further strengthen the *in vitro* results that autophagy flux is more vulnerable than mitochondria to Mn toxicity.

**Fig 5.**
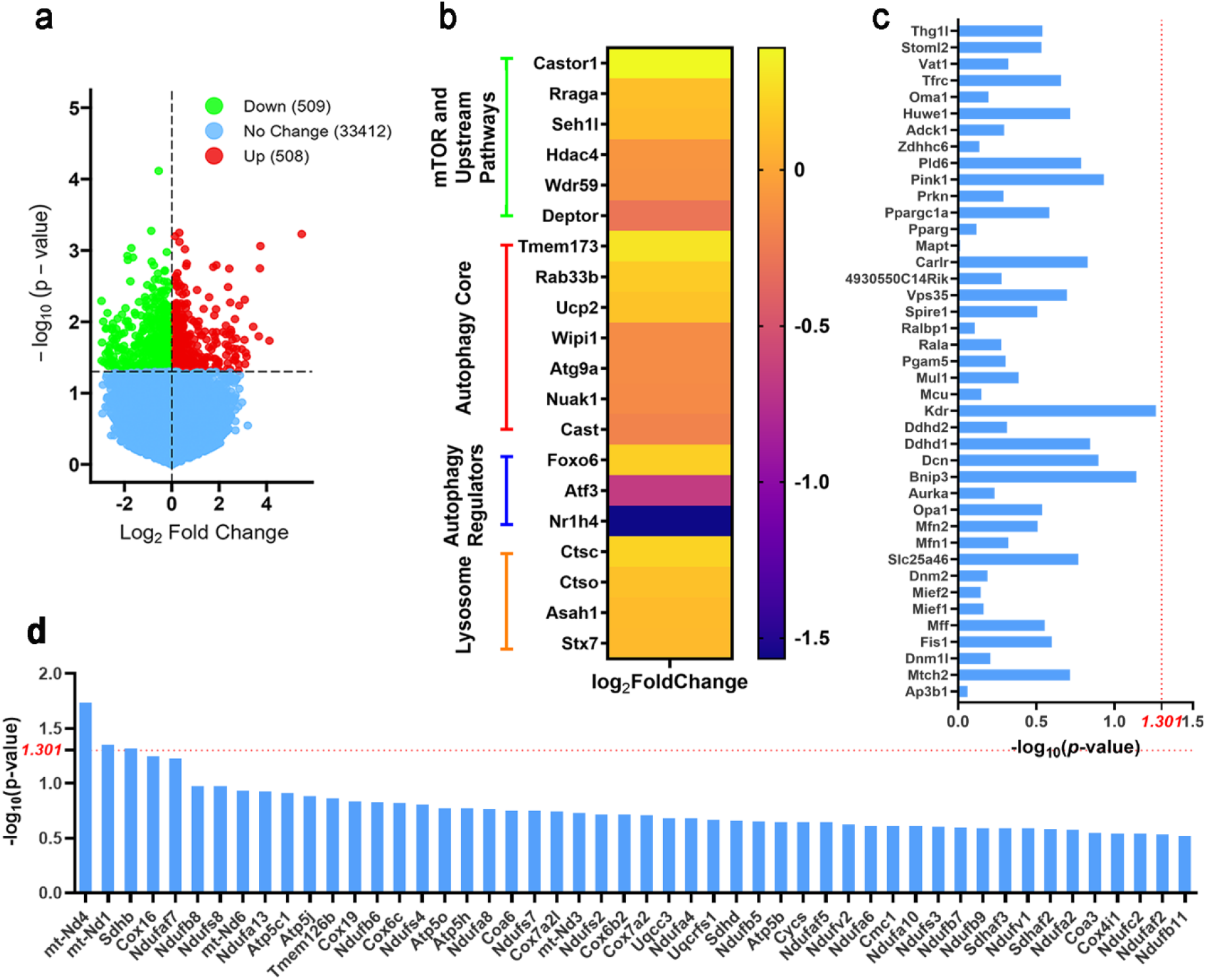
RNAseq reveals Mn impairs autophagy but not mitochondrial pathways *in vivo*. C57/BL6 mice (3–4-month-old) were orally gavaged with MnCl_2_ (15mg/kg/day) or water control for 30 consecutive days before harvesting (n=8 for each group). Total RNA was extracted from the ventral midbrain and processed for RNAseq. **a,** Volcano plot showing the number of differentially expressed genes between the Mn-treated mice and their control littermates (>one-fold change, p<0.05). **b**, Heat map showing differential expressed genes (DEGs) of the autophagy pathways between the Mn-treated mice and the control mice. **c,** Genes related to mitochondrial dynamics were not significantly affected after Mn exposure [-log_10_(p-value)<1.301]. **d**, Genes involved in mitochondrial function show 97.7% (128/131 genes) were not significantly affected in the Mn-treated mice compared to the control group [-log_10_(p-value)<1.301]. For brevity, only the top 50 genes are shown here.

### Mn impairs autophagy flux in nigral dopaminergic neurons

Although our transcriptomic data show impaired autophagy pathways, we further validated autophagy flux at the cellular protein levels using transgenic autophagy reporter mice that express ubiquitously a tandem RFP-EGFP-LC3 fusion protein^39^, the same strategy used to generate the stable autophagy reporter cells used in our in vitro studies (Fig. 1). These mutant mice were treated with the same oral MnCl_2_ regimen as in the RNAseq study. Given the potential role of Mn as a risk factor in PD, we assessed autophagy flux in dopamine (DA) neurons in SNpc (Fig 6a). We also included GABA neurons in the substantia nigra *pars reticulata* (SNpr) as control (Fig 6b). Using confocal microscopy, we captured the red and green fluorescent puncta and quantified the number of autophagosomes and autolysosomes. As seen in Fig. 6c, Mn impaired autophagy in tyrosine hydroxylase (TH, a phenotypic marker for DA neurons) positive neurons as evidenced by an increase in autophagosomes and a decrease in autolysosome. In contrast, Mn did not impair autophagy flux in GAD67 positive-GABA neurons (Fig. 6d). Although the number of autophagosomes was increased, there was no reduction in the autolysosome levels. In fact, there was a marginal, but statistically significant, increase in autolysosomes. Next, we evaluated the conversion rate of autophagosome to autolysosome by calculating the ratio of these two vesicles (Fig. 6e). Mn decreased this conversion rate in DA neurons, suggesting the impairment is mostly impacted by downstream of lysosomal fusion and degradation, rather than the upstream of autophagy induction. This observation is consistent with the *in vitro* data (Fig.1f-i).

**Fig 6.**
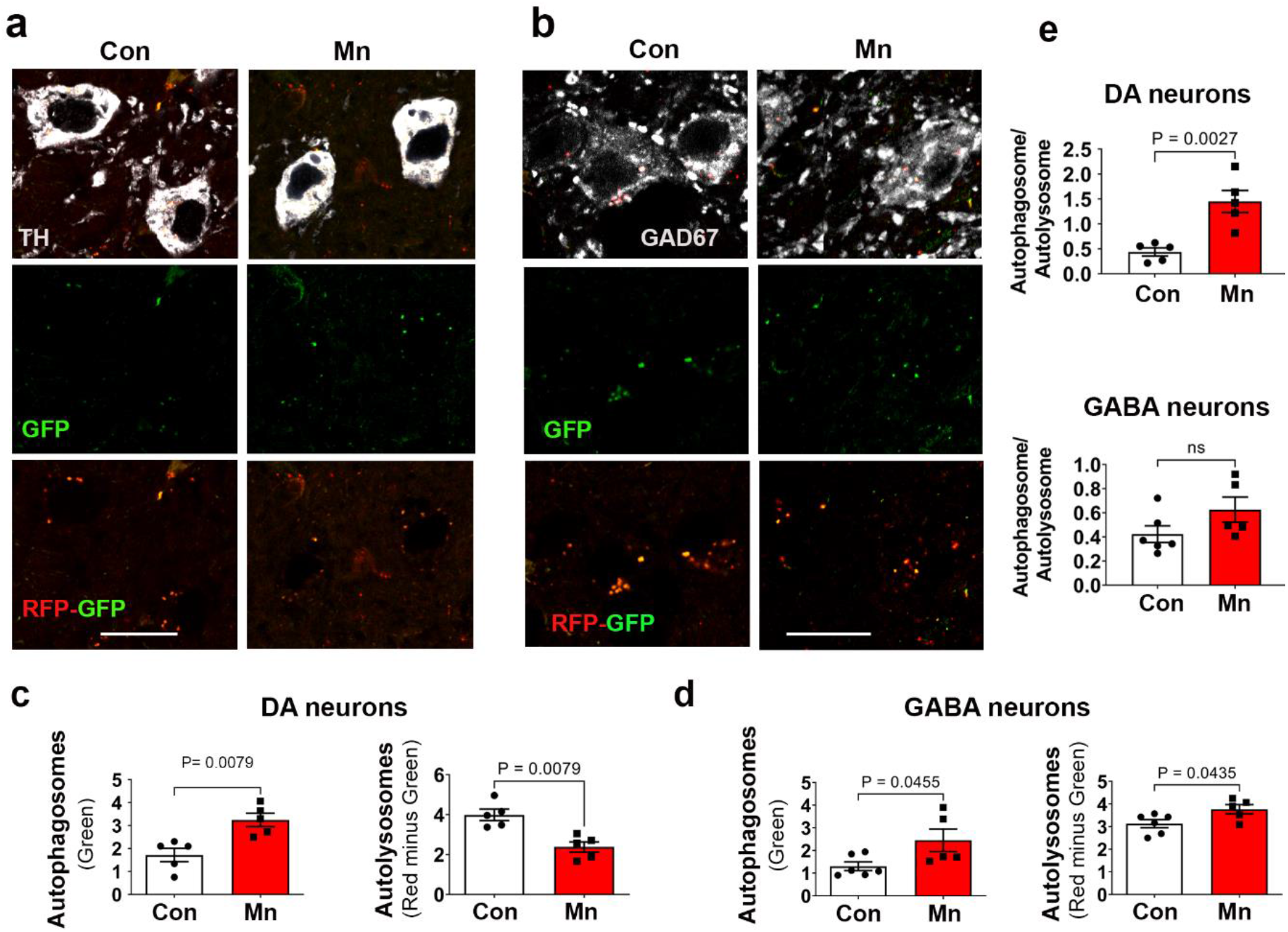
Selective autophagy impairment induced by Mn in the ventral midbrain. Autophagy reporter mice [C57BL/6-Tg(CAG-RFP/EGFP/Map1lc3b)1Hill/J] were treated with MnCl2 (15mg/kg/day) or water through oral gavage once daily for 30 consecutive days, then perfused for immunostaining and confocal imaging of DA neurons **(a**, tyrosine hydroxylase positive) and GABAergic neurons **(b**, GAD67 positive). Autophagic vesicles were quantified for autophagosome and autolysosome and normalized as per 100 µm^2^ for DA neurons **(c)** and GABAergic neurons **(d). e,** The conversion rate of autophagosome to autolysosome was calculated by expressing the ratio of these two types of vesicles. Data represent Mean± SEM, n = 5 per group, with approximately 40-50 neurons analyzed per animal. The number of autophagosomes, autolysosomes, and their ratio were compared using an independent t-test.

### Generation and characterization of Drp1-deficient mice

To evaluate the protective effects of a partial Drp1 deficiency against Mn-induced autophagy in the brain, we generated Drp1-knockout (KO) mice using targeting strategies as illustrated in Fig. 7a. We maintain heterozygous Drp1-KO (Drp1^+/-^) and their wild-type (WT) littermate (Fig 7b) because homozygous disruption of Drp1 is embryonically lethal^40^, and a conditional complete deletion of Drp1 causes dopaminergic cell death in adult mice^41^. Quantitative analysis using qPCR (Fig. 7c) and immunoblotting (Fig. 7d) confirmed an approximately 50% reduction of Drp1 levels in the mutant mice. Physically (Fig. 7e,f) and behaviorally (Fig. 7g), there were no differences between Drp1^+/-^ and WT mice. Stereological cell counting shows a similar number of DA neurons in the SNpc (Fig. 7h). Together, these data indicate that a partial reduction of Drp1 in the heterozygous mice does not negatively impact animal development or DA neuron viability.

**Fig 7.**
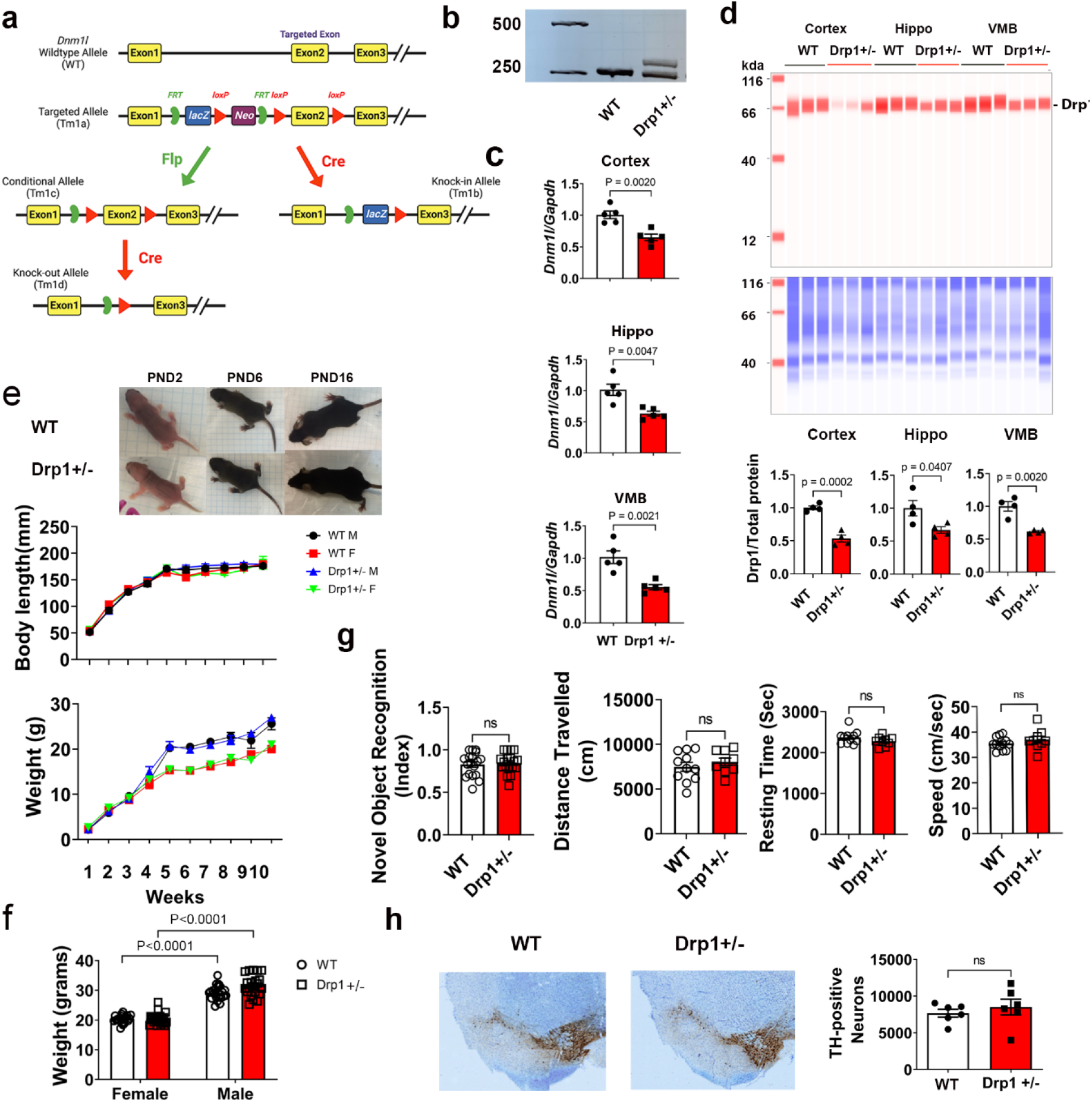
Generation and characterization of Drp1 heterozygous global knockout (Drp1^+/−^) mice. **a**, Schematic diagram illustrating the targeting strategy for generating different types of Drp1-deficient mice using “knockout-first” technology. The KO-first allele is flexible and can produce reporter knockouts, conditional knockouts, and null alleles following exposure to site-specific recombinases Cre and Flp to delete Exon 2 of the *Dnm1l* gene. **b,** representative genotyping results of Drp1^+/−^ versus WT controls. (**c**) qPCR and (**d**) immunoblotting confirmed the reduction of mRNA and protein levels of Drp1 in the Drp1^+/−^ mice in different brain regions (cortex, hippocampus, and ventral midbrain). **e,f,** Weekly measurement showed no differences in body weight and length between Drp1^+/−^ and WT littermates during the developmental stage, n=66 WT (44M & 22F), n=63 Drp1^+/-^ (33M & 30F). **g**, No differences in locomotor activity (n=11 WT, n=9 Drp1^+/-^), as well as learning and memory test (n=18/group) between genotypes. **h**, Stereological counting of DA neurons in the SNpc show no differences between genotypes (data represent Mean± SEM, n=6). Statistical comparison was done using an independent t-test (two groups) and Two-Way ANOVA (two variables).

### Drp1^+/−^ mice is protective against autophagy impairment induced by Mn

Based on the observation that Mn selectively impaired autophagy in nigral DA neurons but not their GABA counterparts (Fig. 6), we used laser microdissection (as illustrated in Fig. 8a) to specifically remove SNpc (rich in DA neurons) and SNpr (rich in GABA neurons) in Drp1^+/-^ mice and their WT littermates treated with oral MnCl_2_. With these tissues, we performed immunoblotting for p62 levels using Jess™ system (ProteinSimple, Inc). Mn significantly increased the p62 protein levels in the nigral DA neurons from WT mice but not in the Drp1^+/−^ mice. However, Mn treatment did not affect p62 protein levels in the nigral GABA neurons from either Drp1^+/−^ mice or WT littermates (Fig.8b-d). To assess whether mitochondrial network was also affected in DA neurons, we quantified mitochondrial morphology in these animals (Fig 8e-g). No differences in mitochondrial network was detectable between genotypes as well as between Mn treatment and vehicle control. Collectively, these results indicate that at a low chronic oral dose, Mn selectively impairs autophagy in nigral DA neurons which are protected by Drp1 inhibition. Furthermore, this protection is independent of the mitochondrial dynamics role of Drp1.

**Fig 8.**
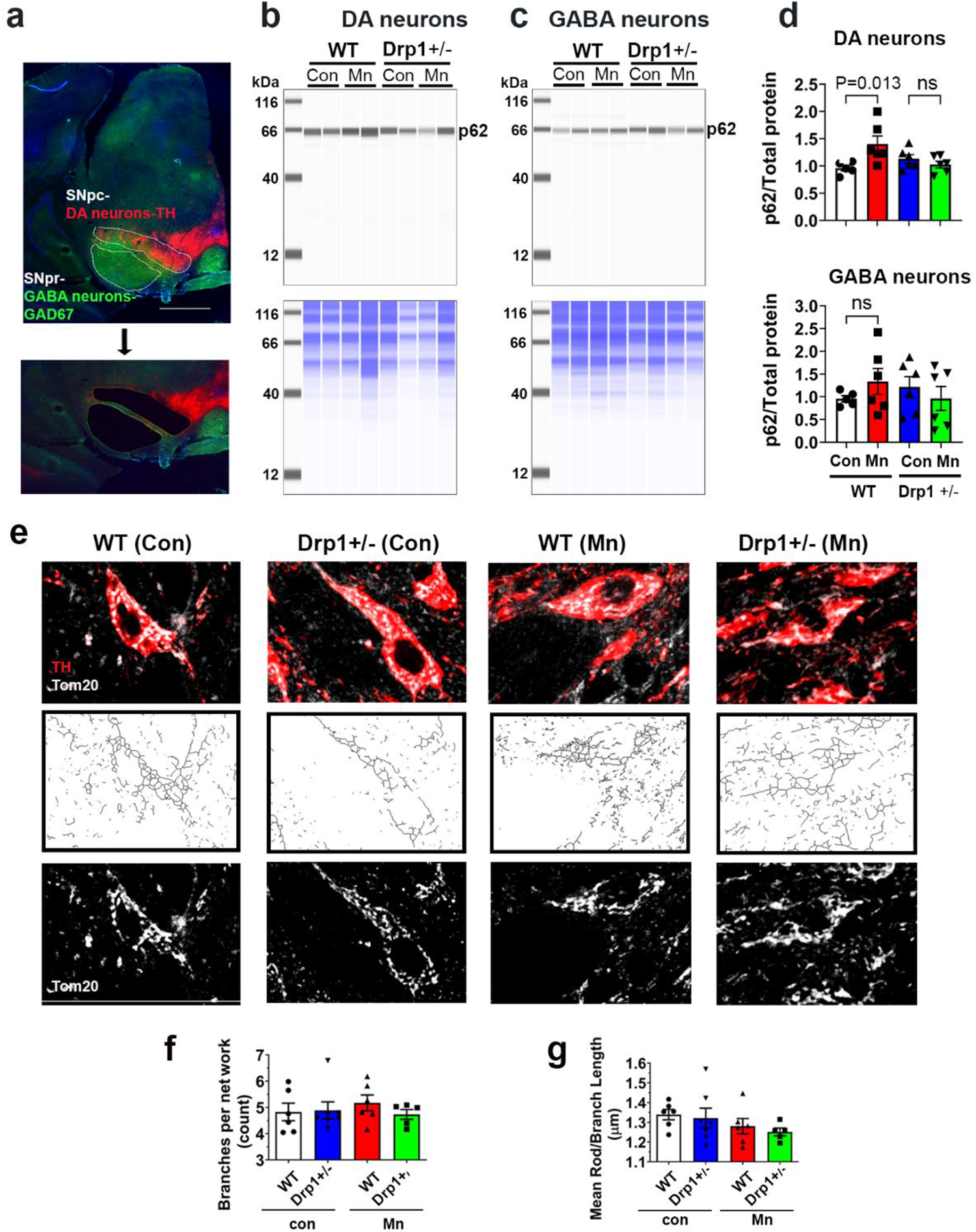
Drp1+/-mice is protective against autophagy impairment induced by Mn. **a**, Representative images of the coronal mouse midbrain section (20 µm) co-immunostained for DA neurons (TH, red) in the SNpc and GABA neurons (GAD67, green) in the SNpr. Both of these brain regions were removed by laser microdissection for immunoblotting. **b,c,** immunoblots of p62 (top panels) in DA neurons **(b)** and GABA neurons **(c)**. Total proteins per lane (bottom panels) were used as loading control. **d,** Quantified levels of p62 were significantly increased in DA neurons, but not in the GABA neurons, of the Mn-treated WT mice. Mn did not significantly increase p62 in DA neurons of the Drp1^+/−^ mice (n=5 for WT control, n=6 for other groups). Data represent mean ± SEM, two-way ANOVA followed by Tukey post-hoc test. **e,** Representative confocal images of mitochondrial morphology of DA neurons after TOM-20 immunostaining (upper panels), then skeletonized (middle panels) for subsequent analysis. Scale bar 20μm. **f,** Various parameters of mitochondrial morphology were quantified using Fiji MiNA plugin. Data represents mean ± SEM (n=6 mice per group), analyzed by one-way ANOVA

## Discussion

Mitochondrial dysfunction is a pathogenic mechanism in neurodegenerative diseases such as AD, PD, HD and ALS^6^. For PD, the discovery of MPTP in the early 1980’s, and the subsequent discoveries of PINK1, parkin being involved in mitochondrial function and quality control^42^, have put the spotlight on the significant role of mitochondria. The observations that PINK1 and parkin affect mitochondrial dynamics has broaden the interest beyond the electron transport chain to include mitochondrial fission, fusion, and movement, not just in PD but also other neurodegenerative diseases such as AD, HD and ALS^6^. Restoring the balance in mitochondrial fission and fusion is one common potential therapeutic approach for these diseases. Blocking Drp1 using both genetic and pharmacological approaches has been shown to be protective in multiple models of neurodegenerative diseases^6^. In addition to improving mitochondrial function and morphology, one striking feature in some of these studies are the observations of reduced protein aggregation of α-synuclein^13, 14^, huntingtin^17^, and amyloid-beta^15, 16^. Although it is possible that the enhanced clearance of these aggregates is due to improved mitochondrial function, it is also possible that blocking Drp1 has a direct impact on improving autophagy to remove misfolded/aggregated proteins.

Autophagy and mitochondria are functionally linked in a bidirectional manner^34^. Autophagy controls mitochondrial quality and number by selectively removing depolarized and damaged mitochondria through a process known as mitophagy. PINK1 and Parkin, the two proteins involved in autosomal recessive PD play a critical role in mitophagy^43^. On the other hand, mitochondria can also control autophagy function. It has been demonstrated that the activity of autophagy is highly dependent on the metabolic state of mitochondria^44, 45^. For this reason, to evaluate whether Drp1 inhibition improves autophagy function independent of improving mitochondrial function, it is critical to have a model without impaired mitochondria since blocking Drp1 itself may improve mitochondrial function, which in turn, improves autophagy. Current genetic models with gene-products linked to neurodegenerative diseases such as PD, AD, HD and ALS have been reported to impair mitochondrial function-either directly or indirectly^46^.

To address the critical question of whether a partial Drp1 inhibition would improve autophagy flux independent of mitochondria, we searched for a neurological disease-relevant model that impairs autophagy without affecting mitochondria. We discovered that at low concentrations, Mn met these criteria. Using two cell culture models (stable autophagy HeLa reporter cells and DA neuronal cells), we performed dose-response studies and revealed that there is a threshold (250µM) that separated the effects of Mn on autophagy flux and mitochondrial function and morphology. Such threshold could be cell type dependent. For example, in primary astrocytes, Mn inhibits both mitochondria and autophagy at 100µM^33, 47^. This observation is consistent with the ability of astrocyte to concentrate Mn up to 50-fold greater than those in neurons through their expression of the transferrin receptors and divalent metal transporter^48^. However, the in vivo significance of such effects of Mn on astrocytes needs to be further validated because despite high accumulation of Mn in astrocyte mitochondria, rats treated with Mn in drinking water (20mg/ml, equivalent to 100mM) for 13 weeks exhibit no ultrastructural damage^49^. To strengthen our in vitro data, we used two mouse models (transgenic autophagy reporter mice and wild-type C57Bl mice). We treated them with a low-dose chronic oral Mn regimen that was previously reported to increase α-synuclein aggregation and transmission via exosome^37^. With these in vivo studies, we discovered that Mn selectively impaired autophagy flux in the SNpc DA neurons but not the nearby GABA neurons in the SNpr. Mitochondrial network structure was not affected by this Mn regimen. RNAseq data further confirmed autophagy pathways were impaired but not mitochondrial related genes. After characterizing the effects of Mn on autophagy and mitochondria, we proceeded to evaluate the effects of a partial Drp1 inhibition on autophagy flux. Due to the concern of potential off-target effects of Drp1 inhibitors, we decided to genetically remove a partial function of Drp1. In cell culture models, we previously validated that siRNA-Drp1 reduced over 70% of Drp1 protein^14^. For in vivo studies, we used our recently generated Drp1^+/-^ mice. In both in vitro and in vivo models, we demonstrated that a partial inhibition of Drp1 protected autophagy flux impairment induced by Mn. In combination, these results demonstrate that a partial Drp1 inhibition improves autophagy flux independent of mitochondrial function.

Autophagy plays a critical role in removing most long-lived and aggregated-prone proteins, including α-synuclein^45, 50–52^. Therefore, a blockade of autophagy flux would increase the accumulation of α-synuclein. Consistent with this mechanism, our data showing that Mn increased the accumulation of proteinase-K resistant α-synuclein. This increase will create a feedback loop^53^ since α-synuclein itself inhibits autophagy function^54^. As a compensatory response to reduce intracellular accumulation of α-synuclein when lysosomal function is blocked, exosome-mediated transfer release of α-synuclein is increased^55–57^. This observation is consistent with the interacting pathways between autophagy flux and amphisome-mediated exosome release^57^. When autophagy flux is blocked, the multi-vesicular body and amphisome pathways are more active to reduce intracellular α-synuclein levels, leading to more exosomal release^57^. Consistent with the inhibitory effects of Mn on autophagy in the present study, a previous study reported that Mn increased the release of exosomes containing oligomeric synuclein^37^. Thus, by improving autophagy function, Drp1 inhibition can reduce synuclein aggregation and spread.

Acute high exposure to Mn causes manganism. Because globus pallidus is the primary brain region affected, these parkinsonian-like symptoms are not responsive to levodopa ^22, 58, 59^. However, low-level long-term exposure to Mn may extend to the entire basal ganglia, including the substantia nigra, as demonstrated experimentally in mice^29^. In human occupational studies, when exposed to welding fumes which contain Mn, welders are more likely to develop PD/parkinsonism^60–62^. Consistently with these previous investigations, a recent study reported that serum exosomes of welders contain higher misfolded α-synuclein^37^. In addition to occupational exposure, Mn in the living environment is also a concern. A high prevalence of parkinsonism has been observed in the province of Brescia, Italy, where residents live in the vicinities of ferroalloys industries operating since the beginning of 1900^27, 63^. In the same province, case-control studies have shown a relationship between PD/parkinsonism with metal exposure and α-synuclein polymorphism^63^. These studies indicate that Mn may be a risk factor for developing PD, especially when interacting with an individual’s genetic makeup^25, 64^. Such interactions have been demonstrated experimentally where Mn is combined with PD-linked genes such as *SNCA*, *parkin, DJ-1,* and *ATP13A2*^24, 48, 65^. Because PD-linked proteins and Mn share pathogenic mechanisms such as mitochondrial dysfunction, neuroinflammation, oxidative stress, and autophagy impaiment^37^, it is not surprising that when combined, their neurotoxic effects are amplified. Our data showing that at low concentrations, Mn impairs autophagy and enhances α-synuclein aggregation, consistent with the proposal that this heavy metal may increase the risk of developing PD.

In summary, the present study provides two major novel mechanisms relevant to neurological disorders. First, we demonstrate that at low concentrations, autophagy is the initial vulnerable target of Mn – not mitochondria as previously reported in studies where high Mn concentrations were used. Second, we provide evidence showing that a partial reduction in Drp1 levels improves autophagy flux independent of mitochondrial function. Given that impaired autophagy and mitochondrial dysfunction are two prominent features of neurodegenerative diseases, the combined protective mechanisms of improving autophagy flux and mitochondrial function conferred by Drp1 inhibition make this protein an even more attractive therapeutic target.

## Acknowledgements

Acknowledgements

## Author contributions

Conceptualized the research: RF, CS, KT

Designed experiments: RF, CS, YL, SS, JP, SL, KT

Performed experiments: RF, CS, YL, SS, JP

Data analysis and interpretation: RF, CS, YL, SS, JP, JR, SL, KT

Wrote the manuscript: RF, CS, YL, KT

Reviewed and edited the manuscript: RF, CS, YL, SS, JP, JR, SL, KT

Contributed new reagents/cell model: SL

Secured funding: KT

## Data availability

RNA-seq will be deposited in the Gene Expression Omnibus (GEO) at the time of publication. Quantitative data that support the findings of this study are available within the paper. All other data that support the findings of this study are available from the corresponding authors on reasonable request.

## Supplementary Materials

Materials and Methods

Supplementary file 1

Supplementary file 2

