## Supplemental Materials and Methods for "A partial Drp1 knockout improves autophagy flux independent of mitochondrial function"

### Cell cultures

HeLa cells with stable expression of mRFP-GFP-LC3 were used as autophagy reporter cells[14]. These cells were grown in Dulbecco's Modified Eagle's Medium (DMEM, Gibco) supplemented with 10% Fetal Bovine Serum (FBS, Gibco), 100U/ml penicillin/ 100µg/ml streptomycin and 100µg/ml G418 (Gibco) at 37 °C in 5% $CO_2$.

Stable rat dopaminergic N27 neuronal cells with inducible expression of human WT α-synuclein were generated using the ecdysone inducible system as previously described[14]. They were grown in RPMI1640 medium, supplemented with 10% FBS, 500µg/ml G418 and 200µg/ml Hygromycin B (Gibco) at 37 °C in 5% $CO_2$. To induce α-synuclein expression, cells were treated with 20µM ponasterone A (an ecdysone-homolog).

### Cytotoxicity using Calcein AM

HeLa cells were plated at a density of $6\times10^3$ cells/well (for 24 h treatments) or $4\times10^3$ cells/well (for 48 h treatments) in 96-well plates, respectively. N27 were plated at $1\times10^4$ cells/well (for 24 h treatments) and $8\times10^3$ cells/well (for 48 h treatments). Cells were treated with varying concentrations (10µM to 2mM) of $MnCl_2$ for 24 or 48 h. Cytotoxicity was assessed using Calcein AM assay (Thermo Fisher Scientific) in accordance with the manufacture's protocol. Fluorescent signal from the Calcein AM dye was measured using a plate reader (Synergy H1, Biotek).

### Cell transfection and siRNA-mediated Drp1 knockdown

Drp1 knockdown in HeLa and N27 cells were performed as previously described[14]. Briefly, pre-designed siRNA against human *DNM1L* (gene encoding Drp1) and rat *Dnm1l* were purchased from Dharmacon Research, Inc. SMARTpool: siGENOME Human *DNM1L* siRNA was used for HeLa cells, while SMARTpool: siGENOME Rat *Dnm1l* siRNA was used for the N27 cells. To monitor LC3 levels in N27 cells, mCherry-hLC3B-pcDNA3.1 plasmid was used. All the transfection was performed using Lipofectamine™ 3000 (Thermo Fisher Scientific) according to the manufacturer's instructions.

**Immunostaining**

*Cultured cells*. Cells were grown on borosilicate cover slips pre-coated with poly-D-lysine in 24-well plates. Prior to immunostaining, cells were fixed with 4% formaldehyde (Thermo Fisher Scientific), in warm cell culture media at 37 °C for 20 min. After washing with 0.1M PBS, cells were incubated with 4% normal goat serum (NGS) (Vector Laboratories), 0.1% TritonX-100 in 0.1M PBS for 1 h at room temperature before being processed for immunostaining. The primary antibodies used were rabbit anti-TOM20 (1:500, cat. no. FL-145, Santa Cruz, discontinued), mouse anti-Drp1(1:500, cat. no. 611113, BD Biosciences, rabbit anti-p62/SQSTM1(1:500, cat. no. PM045, MBL), rabbit anti- α Synuclein Antibody (1:500, cat. no. AB5038, Millipore). Corresponding Alexa Fluor® (350, 488, 586, and 633) conjugate secondary antibodies (Invitrogen) were used at 1:500-1:1000 dilution. Samples were incubated in primary antibodies overnight at 4°C, and in secondaries at room temperature for 1h, with three washes in PBS in between all incubation steps carried out on a orbital shaker. The cover slips were then mounted on to slides using Prolong™ gold anti-fade mount with or without DAPI (Thermo Fisher Scientific).

*Brain sections*. Mice were perfused with 0.9% saline followed by 4% PFA (prepared in 0.1M phosphate buffer), the brains were removed and further fixed with 4% PFA overnight at 4 ºC. Sequential overnight incubations in 15% and 30% sucrose in 0.1 M phosphate buffer were conducted before the brains were then snap frozen in 2-methylbutane at -55 °C. Frozen brains were mounted in optimal cutting temperature (OCT) medium, and cryo-sectioned at 30µm using cryostat (CryoStar NX70, Thermo Fisher Scientific). Coronal sections were collected free-floating in 0.1M TBS. Sections from mid brain were washed in 0.1M TBS before non-specific binding sites were blocked in 0.1M TBS, 4% NGS, 0.1% TritonX-100 for 1 h at room temperature. Primary antibodies diluted in 0.1M TBS, 2% NGS, 0.1% TritonX-100 were then added and incubated overnight at 4°C. Following washing, sections were incubated with Alexa Fluor® secondary antibodies (1:1000, Invitrogen), diluted in 0.1M TBS, 2% NGS, for 1 h at room temperature protected from light. Primary antibodies used were chicken anti-tyrosine hydroxylase (1:2000, cat. no. 76442, Abcam), mouse anti-GAD67 (1:500, cat. no. MAB5406, Millipore), rabbit anti-TOMM20 (1:1000, cat. no. ab186735, Abcam). Following immunostaining with antibodies, sections were stained with 5ug/ml DAPI (cat. no. 62247, Thermo Scientific™) for 10min and washed three times before mounted onto microscope slides, cover glassed with Prolong™ gold anti-fade mount (Thermo Fisher Scientific).

All fluorescent images were captured using Olympus Fluoview (FV1200) confocal microscope with 60x or 100x objectives, lasers 405 nm, 473 nm, 559 nm, 635 nm were set at 5-6% for all channels.

**Quantification of autophagic vesicles**

The analysis of autophagy flux in stable HeLa cells expressing mRFP-EGFP-LC3 was performed as previously described[14]. Red and green fluorescent vesicles in cells were imaged using confocal microscopy with 60x time objective. Autophagosomes and autolysosomes from at least 50 cells per treatment group were quantified using Fiji. For N27 cells, autophagy blockage was assessed through quantification of LC3-mcherry puncta together with the immunostained p62 puncta captured and quantified using the same approach as for HeLa cells.

**Mitochondrial Network Analysis (MiNA)**

To assess the effect of Mn exposure on mitochondrial network/morphology, Mitochondrial Network Analysis (MiNA, a macro toolset of Fiji via utilizing existing ImageJ plug-ins) was performed as described[66]. Briefly, mitochondria were visualized by immunofluorescence using TOM20 (Abcam, ab186735) and imaged with the Olympus Fluoview 1200 confocal. Prior to MiNA analysis, 8-bit gray scale images were binarized and skeletonized following the steps of unsharp mask, enhance local contrast (CLAHE), and median filtering using Fiji. After producing morphological skeletons (2D), the “tagged skeletons were analyzed using MiNA, which divides objects (mitochondria) into two distinct types: individuals (puncta and rods but no branches) and networks (with connected branches). “Mean branches per Network” represents the average number of branches per network. “Mean branch length” calculates the average length of all rods and branches.

**Proteinase K digestion**

To determine the formation of insoluble protein aggregation in N27 cells, proteinase-K digestion was performed prior to immunostaining of α-synuclein as previously described [14]. Following imaging, the number of insoluble aggregates was quantified using Fiji. The number of puncta per cell was determined by counting approximately 30 cells per sample with three independent replicates.

**Mitochondrial respiration assay**

Mitochondrial respiration was assessed using Agilent $XF^e$ 96 Extracellular Flux Analyzer (Agilent Inc) as previously described[14]. Briefly, cells were seeded into XF Cell Culture Microplate and treated with different concentrations of $MnCl_2$ for 24 h. The culture media was then replaced with XF assay media (XF base media supplied with 1mM pyruvate, 2mM glutamax, 2mM glucose and 2mM HEPES, pH 7.2), the plate was then acclimatized in a non-$CO_2$ incubator for 30 min and placed in the Analyzer. A standard XF cell mito stress test with sequential injections of Oligomycin, FCCP and Rotenone/Antimycin was conducted.

**Gel-free immunoblotting**

Gel-free immunoblotting was performed with Jess™ system (ProteinSimple, Bio-Techne) using 12-230 kDa Fluorescence Separation Module (cat. no. SM-FL004 ProteinSimple, Bio-Techne), as per manufacturer's instructions. Cultured cells or brain samples were lysed using 1X RIPA buffer with proteinase inhibitor cocktail (Thermo Fisher Scientific) prior to centrifugation at 13,000*g* (4°C, 15 min) to collect supernatant. After BCA protein quantification (Thermo Fisher Scientific), lysates were diluted in 0.1X sample buffer (included in the separation module) and mixed with 5x Fluorescent Master Mix (PS-ST01EZ, ProteinSimple, Bio-Techne) to a final concentration of 0.5µg/µl. Samples were then boiled at 95 °C for 5 min or 50°C for 30 min (for p62), and 1.5ug of protein were loaded for each sample. The primary antibodies used were rabbit anti-Atg5 (1:100, cat. no. NB110-53818, Novus Biologicals, Bio-Techne), rabbit anti-Drp1 (1:100, cat. no. NB110-55237, Novus Biologicals, Bio-Techne) and rabbit anti-p62 (1:50, cat. no. NBP1-48320, Novus Biologicals, Bio-Techne). Anti-mouse and anti-rabbit secondary antibodies were prepared using corresponding detection modules from ProteinSimple (Bio-Techne) following manufacturer's instructions. For normalization, Total Protein Detection Module (DM-TP01) were used in combination with RePlex™ Module (RP-001, ProteinSimple, Bio-Techne). Data were recorded and analyzed using Compass v6.0.

**Animal models and Mn treatment**

All mice in this study were bred, maintained, and characterized at animal care facility of Florida International University (FIU), animal care and procedures were approved and conducted in accordance with Institutional Animal Care and Use Committee at FIU.C57BL/6-Tg(CAG-RFP/GFP/Map1lc3b)1Hill/J mice were purchased from the Jackson Laboratory (#027139). These transgenic autophagy reporter mice express ubiquitously a tandem RFP-EGFP-LC3 fusion protein under the CAG promoter[67]. Hemizygotes were crossed with C57BL/6J (Jackson Laboratory,#000664) mice to establish and maintain the colony.

*Dnm1l*-deficient mice (Dnm1l$^{tm1b/tm1c\ (KOMP)\ wtsi/Ics}$, Institut Clinique de la Souris, France). We contracted the European Conditional Mouse Mutagenesis Consortium (EUCOMM) to generate Drp1-knockout (KO) mice. These animals were generated using "knockout first" technology[68] in

C57BL/6N embryonic stem cells[69]. As described by the mouse producer, this strategy relies on the identification of a ‘critical’ exon common to all transcript variants that, when deleted, creates a frame-shift mutation. The KO-first allele is flexible and can produce reporter knockouts, conditional knockouts, and null alleles following exposure to site-specific recombinases Cre and Flp (Please refer to the illustration in Fig. 7a). We crossed the ‘knockout-first’ (tm1a) mice, which contain an IRES:*lacZ* trapping cassette and a floxed promoter-driven *neo* cassette inserted into the intron of the *Dnm1l* gene with non-specific cre mouse strain to delete the promoter driven cassette and floxed exon of the *Dnm1l* allele. This generates a ‘global’ *lacZ* tagged allele expressing mouse strain that lacks Drp1 in all cell types (tm1b mouse). As the homozygous disruption of Drp1 is known to be embryonically lethal, we maintain the tm1b strain as a global heterozygous Drp1-KO ($Drp1^{+/-}$) by crossing with C57BL/6 mice. These mice are viable, fertile and do not display any identifiable defects compared to control mice as demonstrated in Fig. 7. Mice were crossed for over 10 generations before Mn treatment was carried out.

*Mn treatment*. 3-month-old mice were orally gavaged with either 15mg/kg/day of $MnCl_2$ ($MnCl_2 \cdot 4H_2O$, cat. no. 221279, Sigma-Aldrich) or water control once daily for 30 consecutive days as previously described[37]. Mice were sacrificed one day after the last treatment. Brains were micro-dissected by regions and snap-frozen for further analysis.

**RNA-Seq and Transcriptomic analysis**

Total RNA was extracted from mouse ventral midbrain using TRIzol (Thermo Fisher Scientific)) as per manufacturer’s instructions. RNA quality evaluation, cDNA library construction and Illumina sequencing were performed by Novogene Corporation Inc. Differential expression analysis was conducted using the DESeq2 R package (1.20.0) and applied Benjamini Hochberg multiple test correction. Genes with adjusted $p$-value $< 0.05$ and absolute $\log_2$(FoldChange) $> 0$ were considered as differentially expressed. The gene enrichment analysis of the differentially expressed genes (DEGs) that are specifically associated with autophagy pathways were performed using the “Autophagy Transcription Gene Toolbox”[38]. Kyoto Encyclopedia of Genes and Genomes (KEGG) pathway analyses were further performed to validate the enrichment of autophagy pathways[70]. The DEGs that are involved in mitochondrial dynamics and mitochondrial function were also compared between the Mn-treated mice and the control mice in reference to “Mouse Genome Informatics” and “Mouse MitoCarta3.0”, respectively[71, 72].

**Quantitative RT-PCR**

For RT-PCR, 1µg of total RNA was converted to complementary DNA (cDNA) using iScript Reverse Transcription Supermix (Bio-Rad). qPCR was conducted on a QuantStudio 6 detection system (Thermo Fisher Scientific) using TaqMan assays and TaqMan Fast Advanced Master Mix (Thermo Fisher Scientific). The Taqman assays employed were *Dnm1l* (mouse, assay ID: Mm01342903_m1, Thermo Fisher Scientific) and *Gapdh* (mouse, assay ID: Mm99999915_g1, Thermo Fisher Scientific). *Gapdh* served as the internal control. The reaction conditions were as follows: 50 °C for 2 min, 95 °C for 2 min and 40 cycles of 95 °C for 1 s and 60 °C for 20 s. Relative quantification was performed using the $2^{-\Delta\Delta CT}$ method.

**Mouse body weight and length measurements**.

The monitoring of mice throughout their development allowed for comparison of these growth parameters through various stages of development. At each measurement, bodyweight was recorded along with the length of each animal. To enable standardized estimation of mouse length; in brief, the mouse was placed on laminated graph paper (5mm x 5mm grid size) and when in a relaxed position a photo was taken directly from above. The images were collated and labelled with mouse ID prior to quantification of the mouse length. Using Fiji, the scale was set for each image using the 5mm grid. The use of this grid allowed accurate scale determination regardless of variance in the height from which the image was taken. A midline was then drawn along the center of the mouse and the length measured along the midline. For weight measurement, mice were habituated into the test room for 10-20 minutes and weighed on the electronic scale with a perforated cover.

**Behavioral Studies**

3-4-month-old Drp1$^{+/-}$ mice and littermate controls were used for all behavioral assessments. For locomotor activity recordings, mice were habituated to the testing room for half an hour before testing. Animals were placed into the activity chamber (Activity Cages Monitor SOF-812, Med Associates, Fairfax, VT) and monitored for 60 minutes using activity monitor software (Activity Monitor version 7.0.5.10 SOF-812, Med Associates). All locomotor activity recordings were performed during the dark cycle (after 19:00 pm) when mice were active. Novel Object Recognition test was used to assess learning and memory in mice. Naïve mice were habituated to the rectangle testing chamber for one hour. Twenty-four hours later, mice were placed back into the same chamber with two identical objects and allowed to explore the objects for 10 minutes before returning to the home cage. One hour later, they were returned to the same chamber with one object replaced by a novel object and allowed to explore the objects for 5 minutes while being recorded using Nodulus software. The recognition index was calculated as a fraction of the time taken to explore the novel object to the total exploration time. Typically, mice with intact learning and memory capacity spend more time exploring novel objects than familiar ones.

**Quantification of nigral DA neurons**

Stereological cell counting was performed as described[73]. Briefly, serial coronal sections (30µM) were collected free-floating and every 4th midbrain section spanning caudal to rostral was incubated with anti-tyrosine hydroxylase (1:4000, cat. no. 657012, Millipore), followed by biotinylated goat anti-rabbit IgG (1:200 dilution; cat. no. BA-1000, Vector Laboratories), and avidin-biotin complex (Vectastain® ABC HRP Kit, Vector Laboratories). Immunoreactivity was visualized using diaminobenzidine (DAB). Total numbers of TH-positive neurons in substantia nigra par compacta were counted stereologically using the optical fractionator (Visiopharm stereological software, Denmark). The settings for the optical dissector probe (dissector height and counting frame size) were set up at 100x objective (Olympus, UPlanApo, NA 1.35, oil iris). A sampling percentage of 30% of the ROI was counted, using a counting frame area of 6400μm (80μm x 80μm). A guard zone of 1μm was utilized at the upper and lower limits of the dissector. The CE values were <0.1 for all animals.

**Immunofluorescence-laser microdissection**

Serial coronal nigral sections (20-μm) were cut on a standard cryostat (CryoStar NX70, Thermo Fisher Scientific) with a clean blade from snap-frozen mouse brain tissue (one hemisphere). Six to eight sections from each brain were mounted on PEN membrane slides (4μm, cat. no. 11600288, Leica), and the unfixed sections were immediately stored at -80°C until immunofluorescence-LMD was performed. The frozen sections were thawed at room temperature for 1 min without drying and immersed immediately in cold acetone for 5 min. After fixation, the sections were rinsed briefly in 1x phosphate-buffered saline (PBS), pH 7.4, and subjected to immunostaining. The sections were initially blocked with 5%NGS for 30 min, followed by incubation with primary antibodies for 1 h and secondary antibodies for 30 min; all steps were performed at room temperature. Sections were briefly rinsed with 1x PBS between each step. After counterstaining with DAPI - for 5 min at room temperature, the sections were dehydrated in graded alcohols (30 seconds each) and air-dried. The primary antibodies used were rabbit anti-tyrosine hydroxylase (1:250, cat. no. 657012, Millipore) and mouse anti-GAD67 (1:50, cat. no. MAB5406, Millipore). The secondary antibodies used were: Alexa Fluor 568 goat anti-rabbit (cat. no. A11011, Thermo Fisher Scientific) and Alexa Fluor 488 goat anti-mouse (cat. no. A11029, Thermo Fisher Scientific). Both secondary antibodies were used at 1:100 dilution. Subsequently, the immunostained DA neurons in the substantia nigra *pars compacta* and GABA neurons in the substantia nigra *pars reticulata* were collected by laser microdissection (LMD6, Leica). The microdissected tissues (2-4mm$^2$) were further lysed in 1xSDS lysis buffer (1% (w/v) SDS, 10mM EDTA, pH 8.0, and 50mM Tris-HCl, pH 8.0) prior to gel-free immunoblotting for p62.

**Statistics**

All values are expressed as mean ± SEM. Differences between means were analyzed using one- or two-way ANOVA with different mouse genotypes, treatment or time as the independent factors. When ANOVA shows significant differences, Tukey's post-hoc test was used for pair-wise comparisons between means. All data sets were subjected to a normality test and an equality of variance test. When these criteria were not met, nonparametric analysis was applied (Kruskal-Wallis ANOVA test with Dunn's post-hoc test). In all analyses, the null hypothesis ($H_0$) or the alternative hypothesis ($H_1$) was accepted with an $\alpha$-error (false-positive) $\leq$ 5% and a $\beta$-error (false-negative) $\leq$ 20%, respectively.

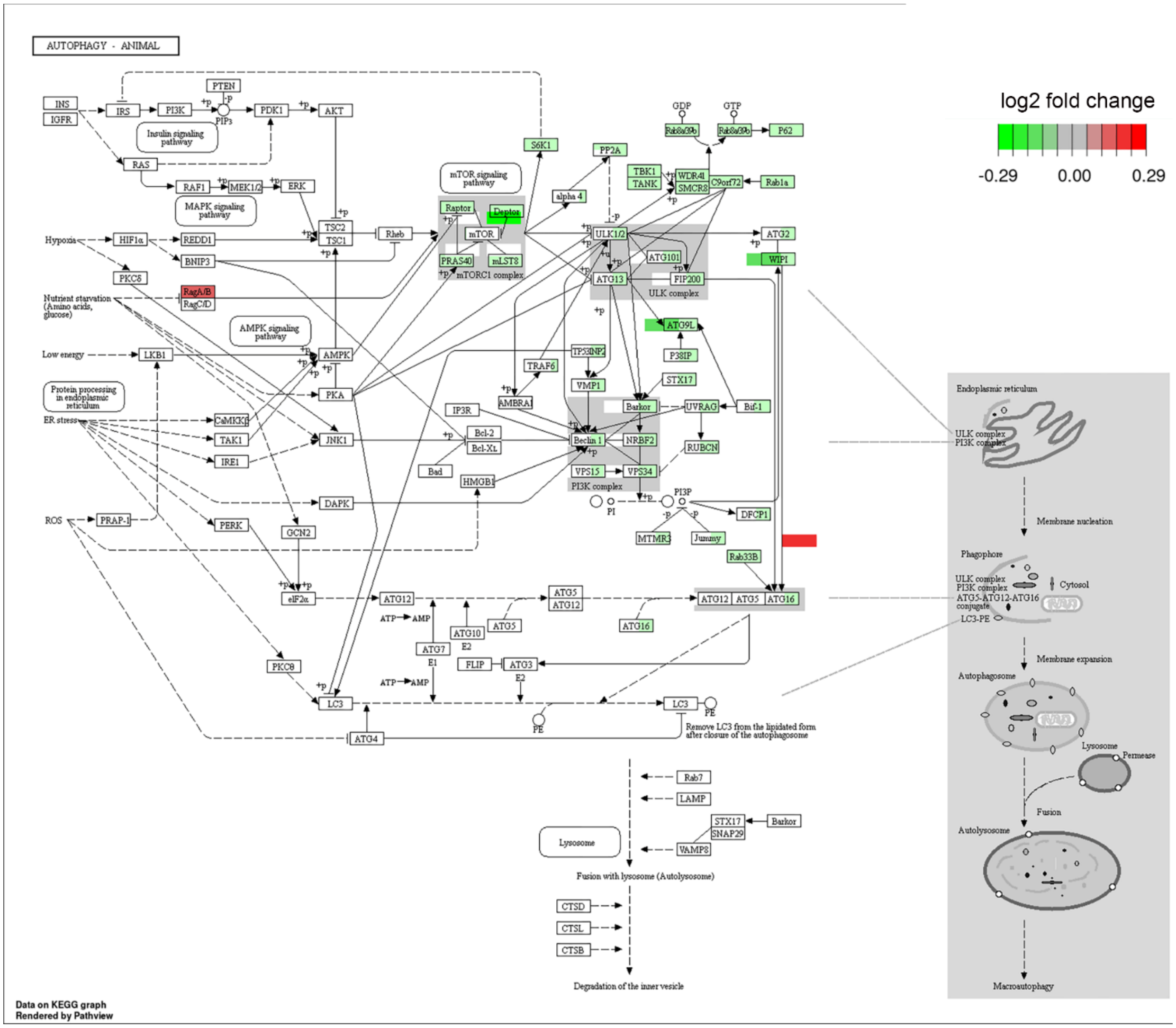


Supplementary Figure 1: KEGG pathways analysis of autophagy associated gene.
